# 4-1BB is a target for immunotherapy in patients with undifferentiated pleomorphic sarcoma

**DOI:** 10.1101/2020.07.10.197293

**Authors:** MJ Melake, HG Smith, D Mansfield, E Davies, MT Dillon, AC Wilkins, EC Patin, M Pedersen, R Buus, AB Miah, SH Zaidi, K Thway, AA Melcher, AJ Hayes, TR Fenton, KJ Harrington, M McLaughlin

## Abstract

Systemic relapse, after treatment of a localised primary tumour with neo-adjuvant radiotherapy and surgery, is the major cause of disease related mortality in patients with sarcoma. As with other cancers, many sarcoma patients derive no benefit from anti-PD-1 treatment. Combining radiotherapy and immunotherapy is under investigation as a means to improve response rates and control metastatic disease. Here, we use a retrospective cohort of sarcoma patients, treated with neoadjuvant radiotherapy, and TCGA data to explore patient stratification for immunotherapy and therapeutic targets of relevance to sarcoma. We show a group of patients with immune-hot undifferentiated pleomorphic sarcoma as one of the highest-ranking candidates for emerging 4-1BB targeting agents. A binary hot/cold classification method indicates 4-1BB-high hot sarcomas share many characteristics with immunotherapy responsive cancers of other pathologies. Hot tumours in sarcoma are however substantially less prevalent. Patient stratification, of intense interest for immunotherapies, is therefore even more important in sarcoma.

---

Soft tissue sarcomas (STS) comprise a heterogeneous group of rare cancers derived from mesenchymal tissue and account for 1% of all adult cancers ^1^. Although capable of occurring at virtually any anatomical site, STS most commonly occur in the limbs and limb girdles, where they are often referred to as extremity soft-tissue sarcomas (ESTS) ^2^. Surgery and radiotherapy delivered pre or postoperatively in patients at high-risk of relapse, remains the cornerstone of management for patients presenting with primary localised ESTS ^3^. Systemic relapses occur in up to a third of patients, accounting for the majority of disease-related mortality ^2,4^. Despite our ability to predict which patients are at highest risk of systemic relapse, the prevention and/or treatment of metastatic sarcoma remains an area of significant unmet need ^5^.

Immunotherapies, most notably immune checkpoint inhibitors (ICIs) targeting the PD-1 (programmed cell death protein 1) and CTLA-4 (cytotoxic T-lymphocyte associated protein axes, have revolutionised the management of patients with historically poor prognoses ^6,7^. However, the initial results from clinical trials with ICIs in patients with metastatic sarcoma were disappointing, with poor response rates in many subtypes ^8–11^. One proffered explanation for this poor response is that sarcomas are ‘immune-cold’ in comparison with more responsive pathologies, often attributed to the low mutational burden in sarcomas ^12–14^. Sarcomas are not a single disease entity but rather a group of cancers with diverse biology and consequently diverse responses to therapy. There is evidence to suggest that patients with undifferentiated pleomorphic sarcomas (UPS) in particular may be more responsive to anti-PD-1 therapy than patients with other sarcoma subtypes ^12,15^. Not only is UPS the most common subtype amongst patients with ESTS, it is also associated with the highest rates of systemic relapse and poorest survival outcomes ^4^. As such, strategies to optimise the response to immunotherapies in this sarcoma subtype are of great interest.

A substantial body of evidence supports the immunostimulatory properties of radiotherapy ^16^. Given its established role in the management of patients with ESTS, the addition of radiotherapy to immunotherapy represents a promising and complementary combination. The immunostimulatory properties of radiation derive from sensing of DNA damage-induced micronuclei by cyclic GMP–AMP synthase (cGAS)–stimulator of interferon genes (STING) ^17–19^. However, the resulting type-I interferon (IFN) response also drives immunosuppressive signalling, such as PD-L1 (programmed death-ligand 1) expression ^20–22^. In preclinical studies, immunotherapy can enhance radiation-induced increases in tumour reactive T cells ^23–25^, and is critical to tumour control outside the radiotherapy field ^26,27^. Clinical findings are beginning to validate these encouraging preclinical data ^28,29^.

In this study, we sought to profile the basal tumour immune environment of patients with UPS prior to treatment with radiotherapy. Our objective was to identify potentially actionable immunotherapy targets that may be treated alone or in combination with radiotherapy. Using historical clinical samples and data from The Cancer Genome Atlas (TCGA), we identify 4-1BB as one such target. 4-1BB is a costimulatory receptor of the TNF receptor family, linked to enhanced survival, cytotoxicity and counteracting exhaustion of T cells, with the natural ligand (CD137L/4-1BBL) expressed on dendritic cells, macrophages and B cells ^30^. Recent studies have however shown 4-1BB to be highly expressed on immunosuppressive tumour infiltrating Tregs ^31,32^. We show levels of 4-1BB to be higher in UPS than other sarcoma subtypes, with 4-1BB expression in UPS amongst the highest in the TCGA dataset. Using a previously published binary hot/cold classification system ^13^, we show levels of 4-1BB and its ligand in hot UPS make it a prime candidate for 4-1BB targeting agents entering clinical trials. We show the immune population landscape of hot sarcoma tumours is in keeping with hot tumours from established immunotherapy responsive cancers. However, hot tumours are significantly less prevalent in sarcoma when compared to other cancers, highlighting the acute need for patient stratification in relation to immunotherapy in these rare tumours.

## Results

### Immune gene expression differs in patients with UPS compared with radiosensitive MLPS

UPS is generally considered to be relatively resistant to both chemo- and radiotherapy. In contrast, myxoid liposarcoma (MLPS) represents a radiosensitive sarcoma when compared with other subtypes ^33^. We sought to compare gene expression between these two subtypes in order to identify markers of response to radiotherapy and potential targets for combination with immunotherapy.

Archival histopathological samples were retrieved from our institution’s tissue bank. These were restricted to patients with a histologically confirmed diagnosis of UPS or MLPS occurring in the extremities who received neoadjuvant radiotherapy prior to surgical resection. Due to molecular similarities between UPS and myxofibrosarcoma (MFS) ^14^, a number of MFS samples were included in non-clustering based analyses. All samples were diagnostic biopsies taken before the start of therapy. Tumour characteristics are outlined in supplementary table 1. UPS patients receiving radiotherapy had a higher rate of progression (Fig. 1a), the shortest time to progression (Fig. 1b), and the highest proportion of progression attributed to distant metastasis (Fig. 1c). These data are in keeping with previous findings from analyses of 556 ESTS patients ^4^.

**Fig. 1.**
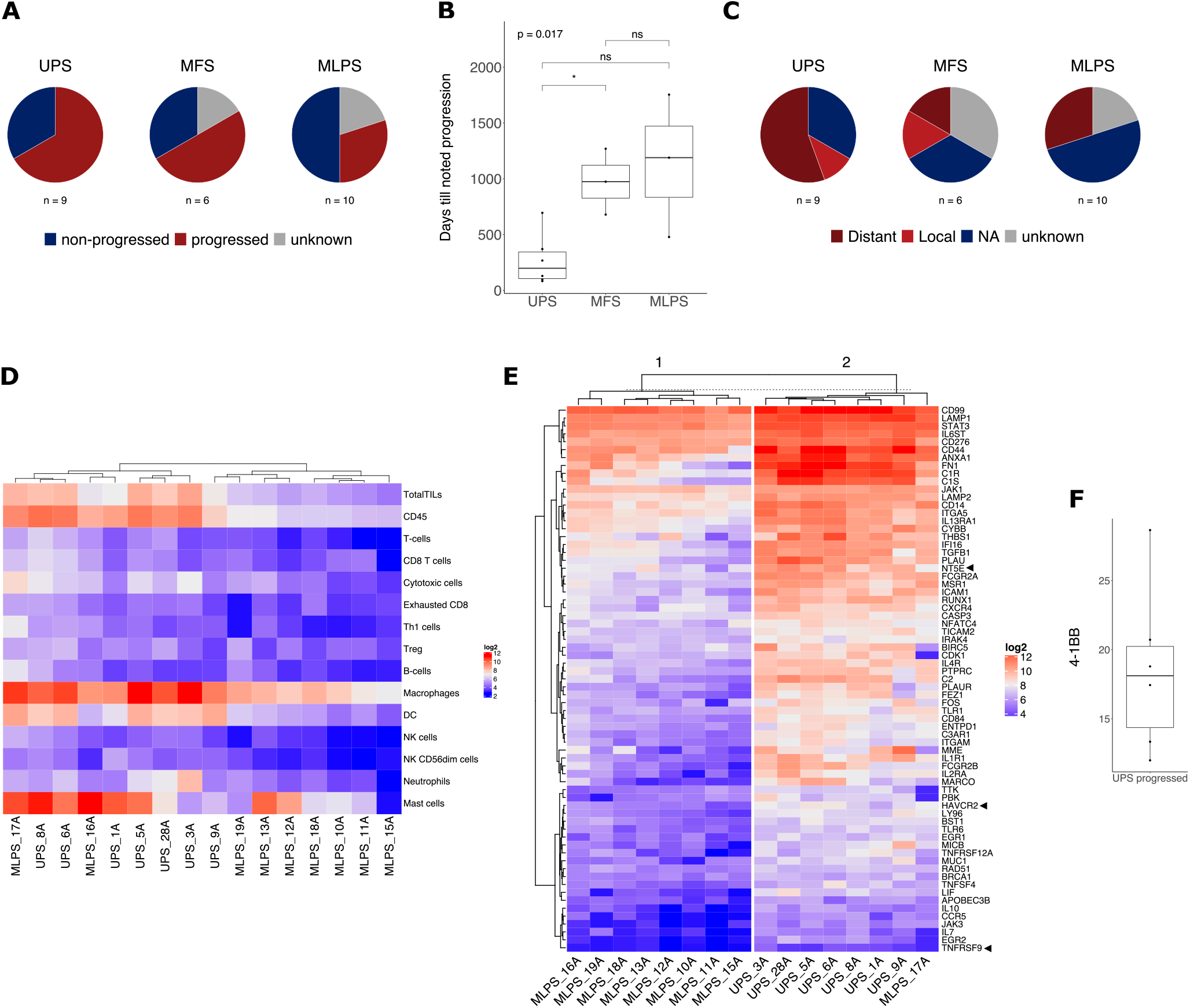
4-1BB (TNFRSF9) is elevated in UPS patients who relapse after radiotherapy. **a.** Progression shown by subtype after neoadjuvant radiotherapy and surgical resection. **b.** Time-to-progression due to local recurrence or metastatic disease. Significance shown t-test, *p<0.05. UPS n=6, MFS n=3, MLPS n=3. **c.** Breakdown of progression by distal, local relapse, unknown, or no progression (NA). Values b-c are in keeping with findings from analyses of 556 ESTS patients 4. **d.** Non-hierarchical clustering analysis of immune cell abundance in MLPS and UPS samples derived from gene expression cell type scoring 34 of NanoString pan-cancer immune panel data. **e.** Differential gene expression of NanoString data comparing UPS and MLPS. Only significantly differentially expressed genes with a fold-change increase of 2 for UPS relative to MLPS are shown.

Tumour areas were outlined by an expert pathologist. After macrodissection, samples with sufficient RNA (Supplementary Figure S1) were analysed using the NanoString pan-cancer immune panel with a custom 30-gene probe set (see methods). Transcript analysis was restricted to a comparison between MLPS and UPS. Immune cell populations were assessed using a validated immune-cell score method ^34^. This indicated higher dendritic cell (DC), macrophage, natural killer (NK) and T-cell populations, as well as CD45 overall, in UPS patient samples compared to MLPS (Fig. 1d). Differential gene expression indicated 69 genes with a statistically significant difference between UPS and MLPS. These are shown grouped by non-hierarchical clustering of all significantly different transcripts (Fig. 1e). Only 1 MLPS sample clustered with UPS. A number of potential immunotherapy targets were differentially expressed, including *HAVCR2* (TIM-3), *NT5E* (CD73) and *TNFRSF9* (4-1BB/CD137, referred to from this point as 4-1BB). After a search of the literature indicated no previous studies profiling 4-1BB in sarcoma, we focused on the potential of 4-1BB as a target for agonists currently under investigation ^30^.

### UPS tumours display colocalization of CD8, B cells, T-regs with high levels of evenly distributed macrophages

Multiplex immunohistochemistry was performed to both corroborate immune-cell scoring (Fig. 1d) and spatially profile the immune microenvironment. The multiplex panel consisted of CD8, CD4, FOXP3, CD68 and CD20 with example images shown (Fig. 2a) alongside quantified population values (Fig. 2b). In keeping with immune-cell scoring data, UPS had higher levels of all populations. MLPS had extremely low levels of all immune cells, except for CD20-positive cells in some biopsies. MFS fell between MLPS and UPS.

**Fig. 2.**
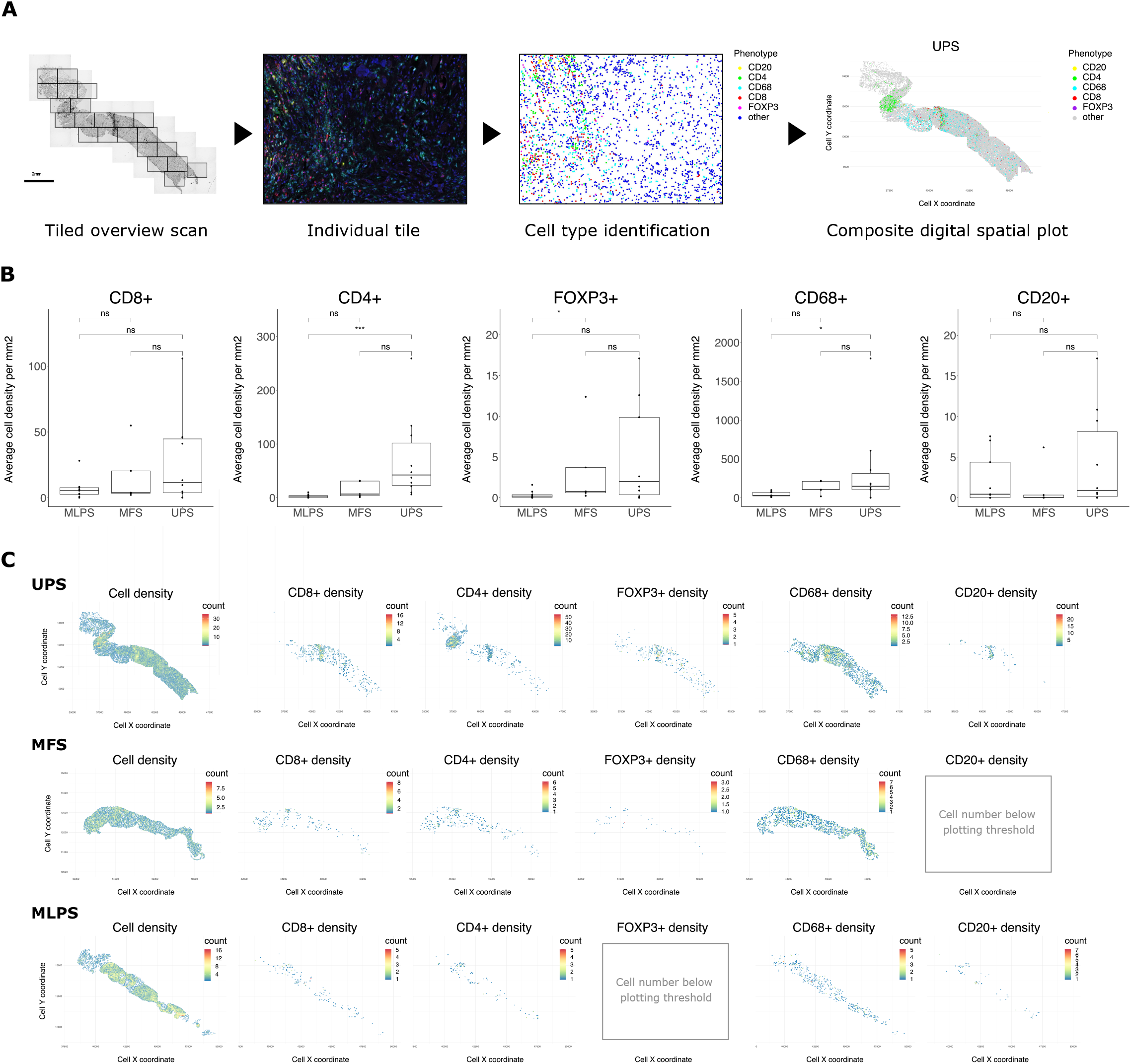
Spatial analysis of immune populations in UPS points to co-localisation of CD8, B cells and Tregs alongside homogeneous macrophage infiltration. Multiplex immunohistochemistry was performed for CD8, CD4, CD20, CD68 and FOXP3 with DAPI as nuclear stain and slides imaged on the Vectra digital pathology imaging system. **a.** A schematic showing tiled images across a biopsy taken at 20x magnification, spectral unmixing and identification of individual cell phenotypes, followed by assembly using coordinate and immune cell phenotype data into a whole biopsy digital spatial composite. **b.** Quantification of immune cell phenotypes was performed for areas of tumour and plotted as cell density per mm2. MLPS n=9, MFS n=5, UPS n=10. Statistics shown, t-test *p<0.05, p***<0.005. **c.** Immune cell phenotypes in digital spatial composites were binned (hexbin) to generate heatmaps of immune cell density across biopsies. Heatmaps for all identified cells, as well as each immune phenotypes alone are shown. Due to differing cell numbers, heatmap scales shown are for each individual plot.

We investigated spatial metrics to put into context any potential co-localisation between populations. Population data from tiled multiplex IHC images were assembled into a digital spatial map of cell populations across biopsies (Supplementary Figure S2). Samples were highly variable in both total immune cell populations and distribution of immune cells within biopsies. To reduce granularity, hexagonal binning was performed on spatial data to generate an immune population density map (Fig. 2c). Due to the number of samples per subtype, as well as the small area provided by biopsies, statistical testing of spatial metrics was not possible. Three spatial distribution patterns were observed. In samples containing areas rich in immune cells, we observed localisation of FOXP3^+^ and CD20^+^ cells with areas of CD8^+^ cells. This was not observed in areas of CD4^+^FOXP3^−^ cells (Fig. 2c). CD68^+^ macrophages were highly prevalent and evenly distributed in a number of samples.

### Analysis of 4-1BB in TCGA data indicates correlation with inflamed and suppressive transcripts

To corroborate our findings that 4-1BB was higher in UPS (Fig. 1) and integrate this observation with co-localization of CD8 cells with suppressive immune populations (Fig. 2), we extended our study to UPS data present in the TCGA dataset. As the analysis of our retrospective cohort (Fig. 1) was performed on pre-radiotherapy diagnostic biopsies, all UPS samples were used in this initial analysis, not just those receiving adjuvant radiotherapy. We first focused on establishing in this larger dataset any correlation between 4-1BB transcript levels and markers of an inflamed or immunosuppressive phenotype. This used a tertile split of UPS samples in the TCGA dataset into high, intermediate and low groups based on 4-1BB transcript levels (Fig. 3a).

**Fig. 3.**
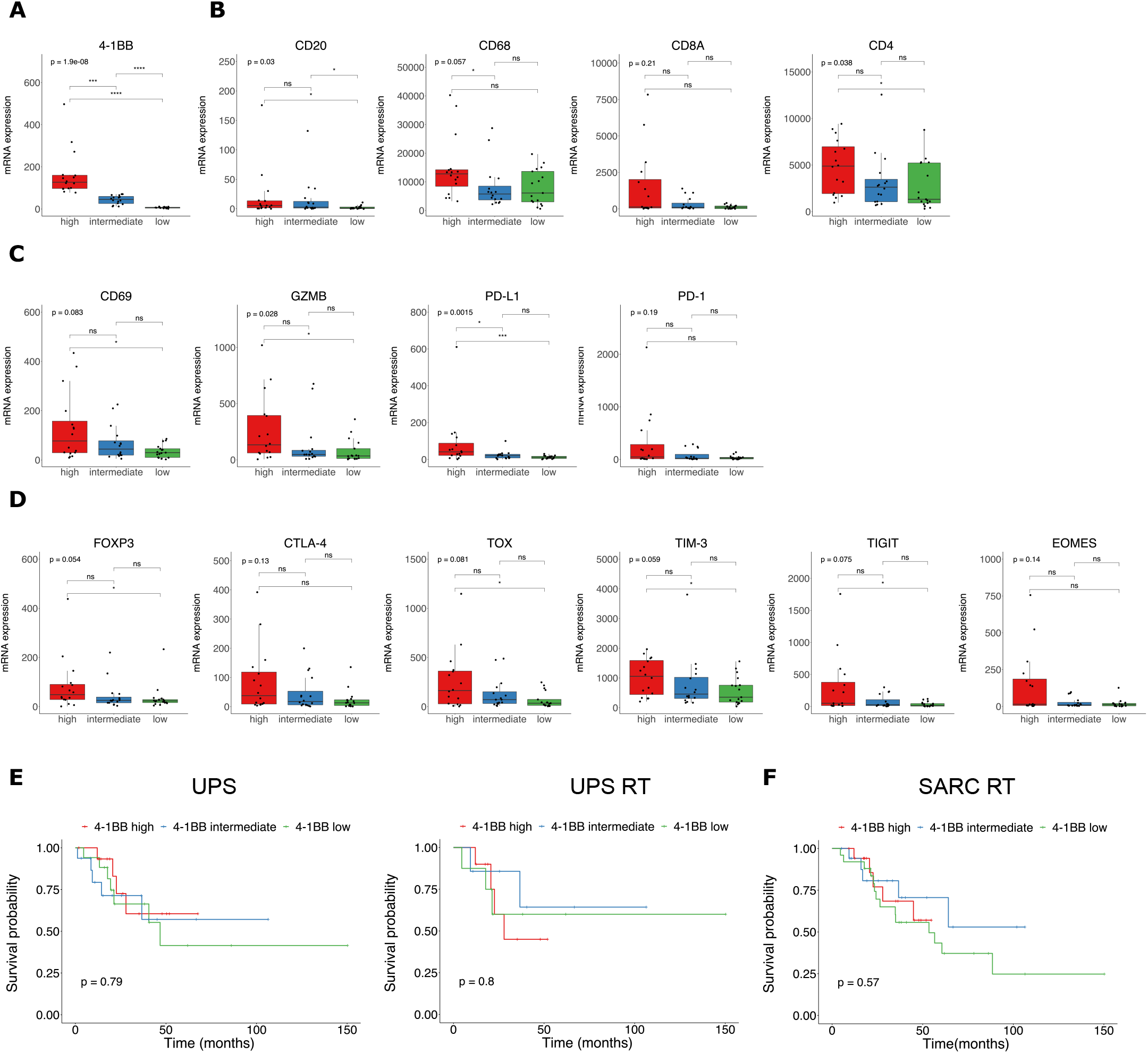
High 4-1BB expression enriches for markers of an inflamed and immunosuppressed phenotype in UPS TCGA data. **a.** A tertile split based on 4-1BB transcript levels was applied to UPS samples in the SARC TCGA dataset. These tertiles are subsequently referred to as 4-1BB-high, -intermediate and -low. Statistics shown are ANOVA, and t-test between groups. For each tertile, mRNA expression was assessed for: **b.** CD20, CD68, CD8A and CD4 transcripts; **c.** cytolytic activity via granzyme B, the activation marker CD69 as well as CD274 (PD-1) and PDCD1 (PD-L1) transcripts; **d.** immune checkpoint, Treg associated transcripts and T cell exhaustion markers. For panels b-d, statistics shown are for kruskal-wallis and wilcox test between individual groups. **e.** Survival probability for all UPS patients by 4-1BB subdivision (left) or restricted to UPS patients receiving adjuvant radiotherapy (right). **f.** Survival probability for all UPS, DDLS, MFS and LMS patients in the TCGA dataset receiving adjuvant radiotherapy by 4-1BB subdivision. For panels e-f, p values calculated by log rank test using ggsurvplot from the survminer package.

Mirroring the spatial profiling performed in figure 2, we looked at *CD20*, *CD68*, *CD8A* and *CD4* in TCGA data for UPS split by 4-1BB status (Fig. 3b). This indicated that within UPS, 4-1BB-high correlated to increased *CD8A* and *CD20*, but less clear trends for *CD4* and *CD68*. Looking at transcripts linked to cytolytic activity in more detail, there was a significant difference in *GZMB* and the activation marker *CD69*, alongside evidence of increased *PDCD1* (PD-1) and *CD274* (PD-L1) transcripts in the 4-1BB-high group (Fig. 3c). The most consistent increase in transcripts linked to 4-1BB levels was observed for immune checkpoints and Treg-associated transcripts (*HAVCR2*/TIM-3*, TIGIT, CTLA-4, FOXP3*) as well as *TOX* and *EOMES* (Fig. 3d), associated with T cell exhaustion.

To determine if 4-1BB transcript levels were linked to patient outcomes, we assessed survival in 4-1BB-high, intermediate and low groups for either all UPS patients in the TCGA dataset or only those receiving adjuvant radiotherapy (Fig. 3e). 4-1BB was not a strong prognostic marker in patients with sarcoma treated with radiotherapy. This is in line with data for PD-L1 where it is not a prognostic marker in NSCLC patients treated with chemotherapy ^35^, but is linked to response rates to PD-1 blockade ^36^. The 4-1BB tertile cut-offs from UPS were applied to all sarcoma subtypes in the TCGA dataset and survival probability plotted for those receiving radiotherapy (Fig. 3f). This extended analysis also indicated no link between 4-1BB and survival.

To summarise, 4-1BB is significantly elevated in UPS versus MLPS in our retrospective cohort. Assessment of immune populations in our retrospective cohort, and subsequent TCGA analyses, indicates an inflamed, but immunosuppressive, environment in UPS patients with high 4-1BB levels.

### 4-1BB expression in UPS ranks amongst the highest in TCGA dataset and immune-hot UPS tumours have similar CD8 levels to immunotherapy-responsive cancers

Our analysis up until this point had been restricted to sarcoma. We were concerned that 4-1BB in UPS, whilst high amongst sarcoma subtypes, may not be high relative to other highly inflamed tumour types. We sought to extend our analysis of 4-1BB in UPS to compare levels across the TCGA dataset. Analysis of 4-1BB transcripts across LMS (leiomyosarcoma), DDLS (dedifferentiated liposarcoma), MFS and UPS in the TCGA dataset indicates UPS contains higher levels than other subtypes (Fig. 4a). Plotted against other cancer types in the TCGA dataset, 4-1BB transcripts in UPS were amongst the highest (Fig. 4b). Non-ligand blocking 4-1BB agonists, such as urelumab, have been shown to promote ligand-dependent receptor clustering ^37^. For this reason, we also assessed 4-1BBL (CD137L/*TNFSF9*) levels across sarcoma subtypes (Fig. 4c) and against other cancer types in the TCGA dataset (Fig. 4d). 4-1BBL transcripts mirrored the findings of 4-1BB, with some of the highest levels detected in UPS and MFS.

**Fig. 4.**
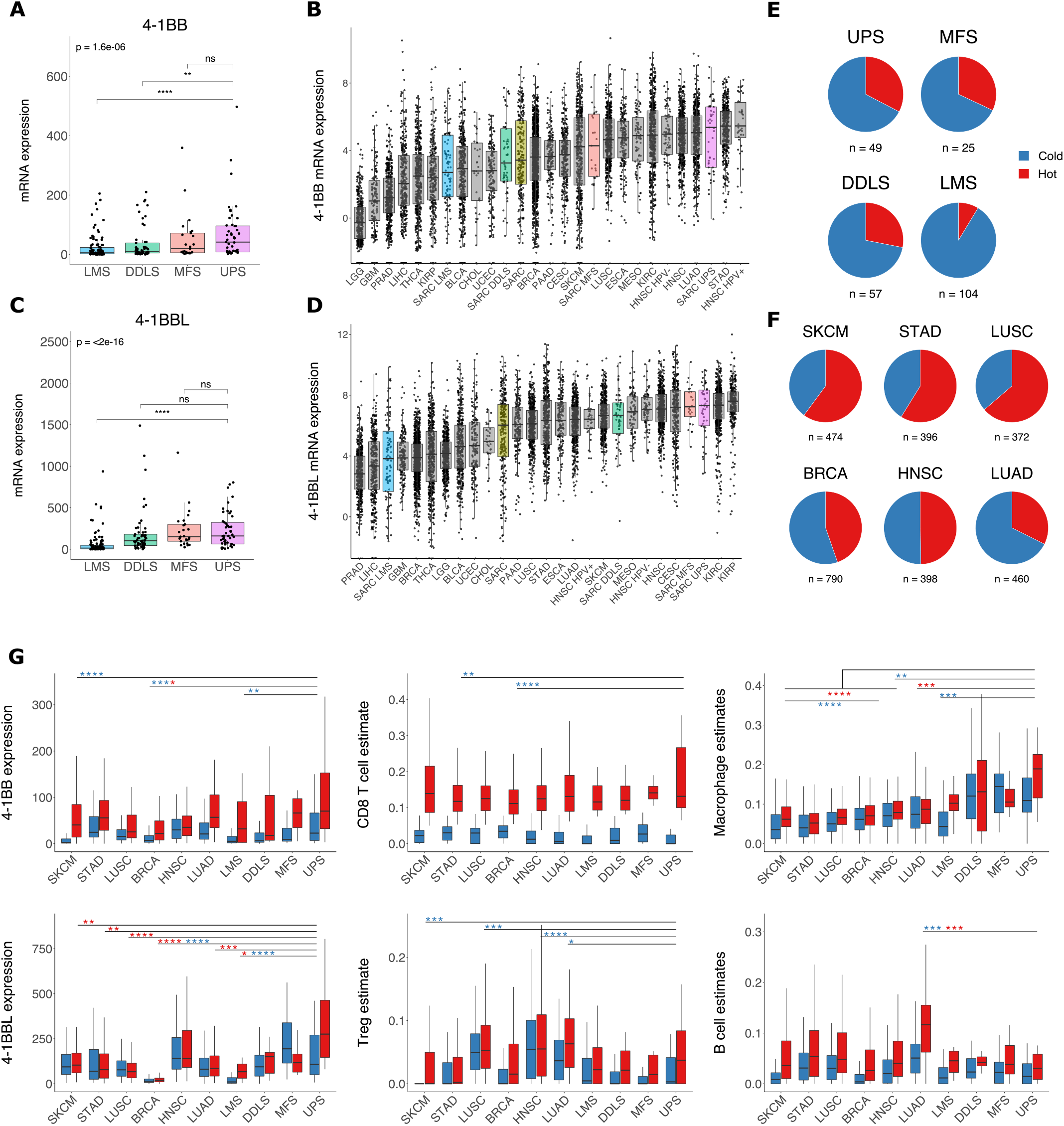
Levels of 4-1BB and 4-1BBL in UPS are amongst the highest in the TCGA dataset. **a.** 4-1BB expression by sarcoma subtype in the TCGA dataset. Statistics shown are kruskal-wallis test, and wilcoxon test between individual groups. **b.** 4-1BB expression in sarcoma subtypes compared to a range of immune-cold and established immunotherapy-responsive cancers in the TCGA dataset. **c.** 4-1BBL expression by sarcoma subtype and **d.** compared to other cancers. **e.** MethylCIBERSORT-derived binary hot/cold immune classification status of sarcoma subtypes and **f.** established immunotherapy-responsive cancers in TCGA. **g.** A comparison between 4-1BB and 4-1BBL expression in hot vs cold sarcoma subtypes compared to established immunotherapy-responsive cancers as well as MethylCIBERSORT population estimates for CD8, Treg, B cell and CD14 macrophages/monocytes. Statistical comparisons in g are between immune-hot or immune-cold UPS and the corresponding hot (red asterisks) or cold group (blue asterisks) as indicated. Pairwise wilcox test with Bonferroni correction, *p<0.05, **p<0.01, ***0.001, ****p<0.0001.

We initially adopted a tertile split of 4-1BB. To further validate our findings, we sought an independent immune classification approach that would allow a ‘like-for like’ comparison between sarcoma and immunotherapy-responsive cancer types. We used a published method of binary classification (immune-hot or immune-cold) based on MethylCIBERSORT-derived estimates of cell abundances in mixed tumour populations ^13^. This binary hot/cold classification indicated UPS, MFS and DDLS possessed similar levels of immune-hot tumours, with LMS substantially colder (Fig. 4e). Sample numbers per subtype classified as hot/cold were: UPS, 16/33; MFS, 8/17; DDLS, 16/41; LMS 9/95. 4-1BB levels do not simply track either the prevalence of hot tumours or the expression of *CTLA4*, *PDCD1* or *CD274* (Supplementary Figure S3). Immune-hot sarcomas were less frequent than other immune-hot cancer types (Fig. 4f), but still comprised over a quarter of UPS, MFS and DDLS. These findings are consistent with a recent study on sarcoma immune classes using a similar gene-expression-based TME deconvolution tool, MCP-counter ^12^.

To establish how hot/cold UPS and other sarcoma subtypes are compared to immune-responsive tumour types ^6,7,29,38^ we compared 4-1BB/4-1BBL and immune population estimates to those in lung, head and neck, melanoma and breast cancer using MethylCIBERSORT-derived population estimates ^13^. 4-1BB transcripts were highest in immune-hot tumors across all sarcoma subtypes, with significant differences observed in MFS and UPS (Supplementary Figure S4). The highest average values for 4-1BB and 4-1BBL transcripts were identified in hot-UPS (Fig. 4g). Whilst 4-1BBL levels appeared higher in hot compared to cold UPS, this was not in keeping other sarcoma subtypes or cancers where little variation was observed between hot and cold (Fig. 4g). Additional analysis of 4-1BB versus 4-1BBL expression does not clearly point to any correlative link in any cancer type (Supplementary S5). Therefore, 4-1BBL expression in UPS is high amongst cancers in the TCGA data, but does not appear to be clearly linked positively or negatively to 4-1BB levels.

In tumours classified as hot, CD8 estimates were similar across all sarcoma subtypes and immunotherapy-responsive cancers (Fig. 4g). Treg estimates were higher in hot sarcoma subtypes and many ICI-responsive cancers, but more variable than for CD8 estimates (Fig. 4g). CD4 effector and NK cell estimates were consistently higher in cold vs hot classifications (Supplementary Figure S6). High CD14 monocyte/macrophage estimates in hot UPS, as well as high levels of NK cell estimates in cold-UPS (Fig. 4g, Supplementary Figure S6) stood out compared to immunotherapy-responsive cancers.

To summarise, immune-hot sarcomas across all subtypes have similar estimated levels of CD8 T-cells compared to established immunotherapy-responsive cancers, and show similar trends for CD4 effector, Treg and B cell estimates. 4-1BBL transcripts are high in UPS compared to other cancers. Hot UPS tumours have some of the highest levels of 4-1BB not only when compared with other sarcoma subtypes, but also compared with hot tumours from immunotherapy responsive cancer types.

### 4-1BB-high levels correlate to CD8, Treg and B cell estimates but with significant heterogeneity of immunosuppressive populations

We returned to the previous tertile split of 4-1BB transcript levels in conjunction with MethylCIBERSORT-derived population estimates to assess correlation between immune populations in 4-1BB-high vs -low groups across all sarcoma subtypes (Fig. 5a). Positive-correlation was observed between CD8, Treg and CD19 estimates in the 4-1BB-high group that was not observed for low 4-1BB. This correlation was consistent across all sarcoma subtypes, including LMS. No clear difference in correlation was observed for CD14 monocytes/macrophages or neutrophils based on 4-1BB levels (Fig. 5a).

**Fig. 5.**
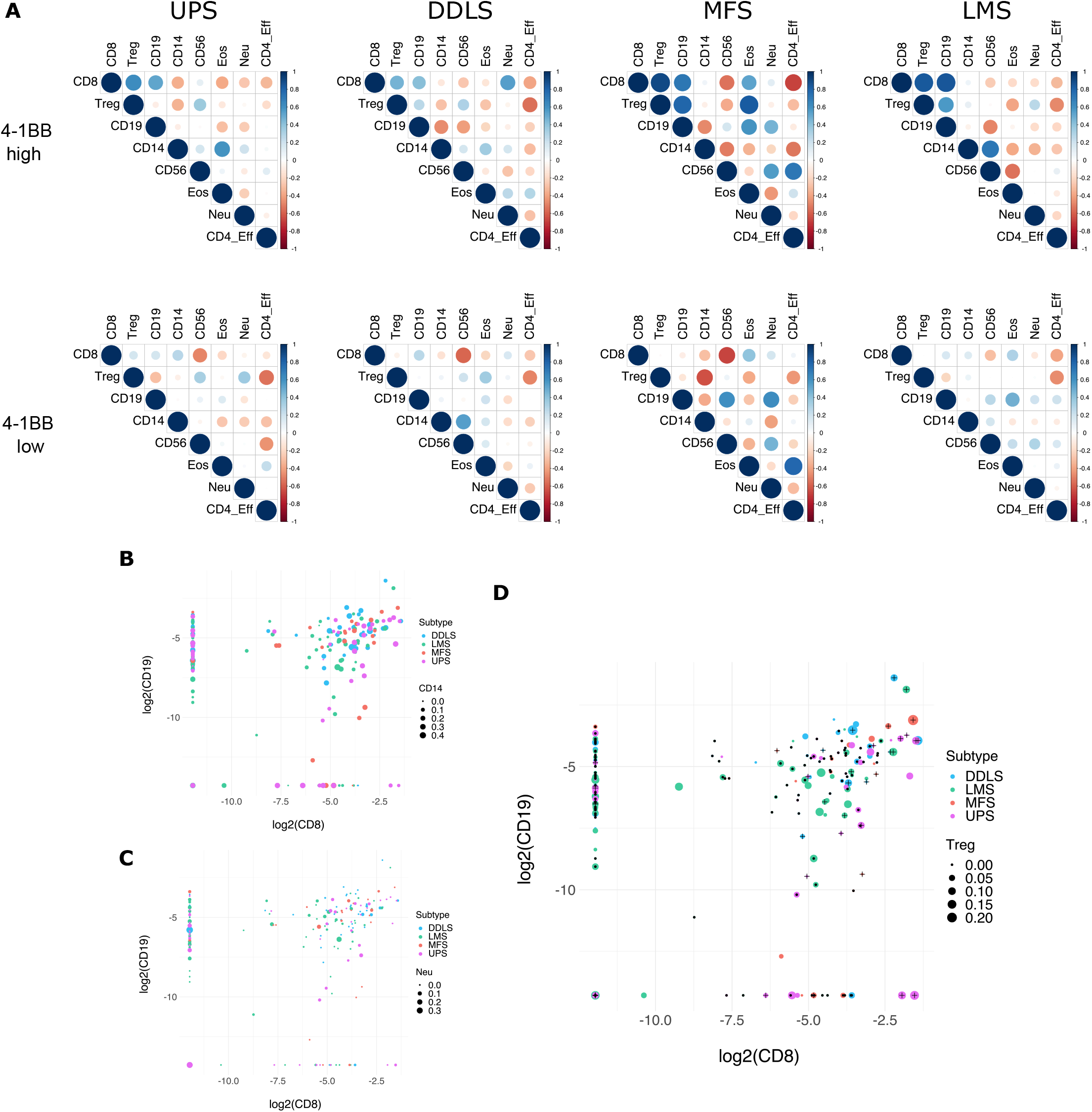
High 4-1BB correlates to B cells, CD8 T-cells and Tregs but immune population co-localisation is highly heterogeneous when viewed on a per sample basis. **a.** Correlation analyses between TCGA MethylCIBERSORT-derived immune cell population estimates was performed in 4-1BB-high and 4-1BB-low tertile groups in TCGA data. Immune population estimates are indicated on the x and y axes. The size of dots at the point of intersection are Pearson’s correlation values with the size of the dot illustrating the extent of correlation (larger equals greater correlation values). Positive correlation shown in blue, negative in red for each subtype. Heterogeneity of potentially suppressive CD14 monocyte/macrophages (**b.**), Neutrophils (**c.**), and Treg populations (**d.**) co-occurring with CD8highCD19high population estimates were assessed. Log2 transformed cell population estimates were plotted for CD8 vs CD19 for all sarcoma subtypes. On top of this xy data, the estimated abundance of CD14, Neutrophils or Treg populations in each tumour is shown by the size of the data point with the legend indicating the estimated values corresponding to the point sizes shown. CD14 estimates are shown to be homogeneously high. Neutrophil estimates are heterogeneous, with a small number of high population values in CD19highCD8high samples. In addition to plotting estimated values for Tregs by point size, samples were also labelled as 4-1BB-high (+), 4-1BB-intermediate (no markings) or 4-1BB-low (dot). Treg population levels are shown to be highly heterogenous, with high and intermediate 4-1BB levels visible on both Treglow and Treghigh datapoints for CD8highCD19high samples.

Monocytes/macrophages (as mononuclear-MDSCs), Tregs and neutrophils (as polymorphonuclear-MDSCs) have all been linked to radiotherapy resistance, either basally, or due to increases shown in preclinical models after radiotherapy ^20,39,40^. We wanted to look at these populations at a more granular level across all four sarcoma subtypes. Since B cells and CD8+ cytotoxic T-cells (CTLs) were linked to response to pembrolizumab in the SARC028 trial ^12^, We interrogated CD14+, neutrophil or Treg population estimates in tumours with higher estimates for B-cells and CTLs to understand better their heterogeneity specifically in this dual-positive group (Fig. 5b-d). These fell into three categories. CD14+ monocytes/macrophages, showing no clear correlation based on 4-1BB levels (Fig. 5a), were relatively homogeneous across all four sarcoma samples (Fig. 5b). In conjunction with spatial data (Fig. 2c) and the higher macrophage/CD14 levels in UPS (Fig 1d, Fig. 4g), monocyte/macrophage populations are a ubiquitously co-localising immune population in hot UPS. Any suppressive role is, therefore, likely to be linked more strongly to other factors such as PD-L1 expression and polarity. Neutrophils, also showing no clear correlation based on 4-1BB, showed some signs of heterogeneity in the dual-positive B-cell and CTL high group (Fig. 5c). Tregs, which did show signs of correlation based on 4-1BB levels, were highly heterogeneous in the dual-positive B-cell and CTL high group (Fig. 5d). We layered the 4-1BB tertile status (dot for low, plus sign for high, intermediate unlabelled) on this final plot (Fig. 5d). In keeping with publications showing high 4-1BB on intratumoural Tregs, many CTL^high^B-cell^high^Treg^high^ samples fell into the 4-1BB-high or -intermediate grouping, with CTL^high^B-cell^high^Treg^low^ samples in the 4-1BB-low group. However, many CTL^high^Treg^low^ samples also fell within the 4-1BB-high and intermediate groups.

These data suggest that, even when restricted to tumours containing both B cell and CD8+ CTL populations, categorising suppressive populations relevant to radiotherapy and immunotherapy in sarcoma is likely essential due to the heterogeneity observed. As 4-1BB has a dichotomous role, expressed on CTLs and Tregs, this could be doubly important when selecting between 4-1BB agonism or depleting antibodies targeting Tregs.

### Hot UPS tumours express elevated T-cell chemoattractants, ISGs and antigen processing

Having looked at immune populations linked to suppressive signalling in response to radiotherapy, we extended our analysis to look at cytokine signalling of potential relevance to radiotherapy. We returned to the hot/cold classification within sarcoma subtypes to analyse differences in T-cell chemoattractants *CXCL9, CXCL10* and *CXCL16*, as well as *CCL2* and *CCL5* due to links to Tregs and MDSCs ^39^ (Fig. 6a). Despite similar levels of CTL estimates across subtypes, the differential between T-cell chemoattractants was significantly more robust in hot vs cold UPS and MFS compared to DDLS and LMS. The most pronounced hot-cold differential for *CCL2* and *CCL5* was observed for UPS.

**Fig. 6.**
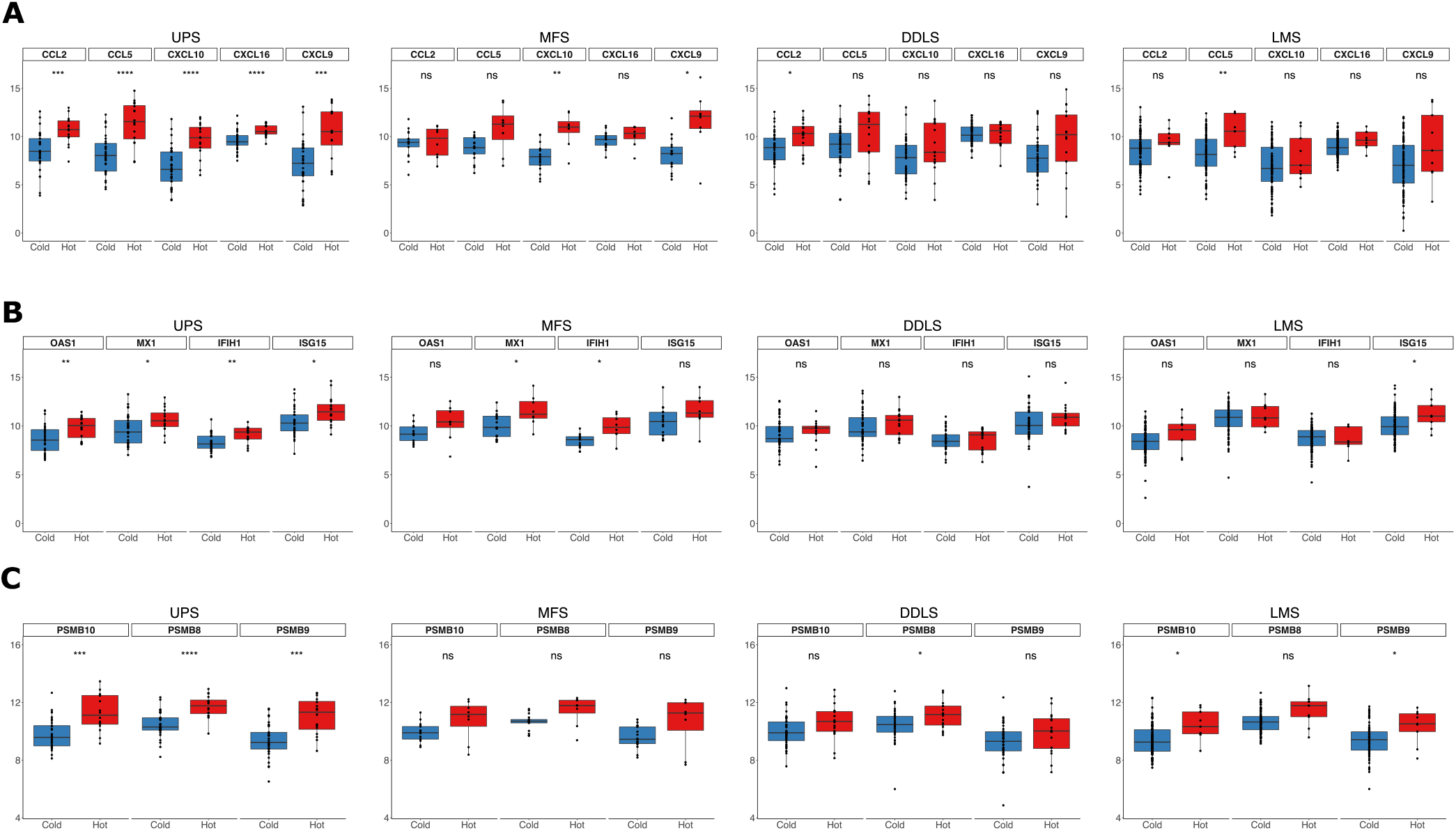
T-cell chemoattractants, interferon-stimulated genes and antigen processing are higher in immune-hot versus immune-cold UPS. **a.** T-cell chemoattractants (CXCL9, CXCL10, CXCL16), and those linked to immunosuppressive Tregs and MDSCs (CCL2, CCL5), were assessed across subtypes based on hot/cold classification of TCGA data. **b.** The expression of interferon-stimulated genes linked to DNA damage resistance (OAS1, MX1, IFIH1, ISG15) was compared between hot and cold tumours by subtype. **c.** Immunoproteasome subunit expression was compared across subtypes between hot and cold classified tumours. Significance shown t-test, *p<0.05, **p<0.01, ***0.001, ****p<0.0001.

Type-I IFN signalling on DCs after radiotherapy has been shown to be therapeutically critical to generation of anti-tumour immunity ^41^. Paradoxically, the presence of an elevated basal interferon-related gene signature has also been linked to resistance to DNA damaging agents clinically ^42^ and radiotherapy-ICI combinations preclinically ^43^. We observed a similar pattern of expression of the interferon-stimulated genes (ISGs) *OAS1*, *MX1*, *IFIH1* and *ISG15* (Fig. 6b) as observed for chemoattractants (Fig. 6a). Statistical significance was only observed between hot and cold UPS or MFS (Fig. 6b). However, there was much greater overlap in values between hot and cold tumours than was observed for chemoattractants. These data are of relevance to radiotherapy in sarcoma due to concerns that high levels of T cell chemoattractants and ISGs could result in resistance to DNA damaging agents, as described for other cancers. Whilst significant, the broad overlap indicates ISG expression does not break down strictly along the lines of hot tumours being more clearly at risk from this reported mechanism of DNA damage resistance.

High clonal neoantigen burden has been shown to correlate to increased expression of CD8, CXCL9/10 and antigen presentation genes in lung adenocarcinoma ^44^. Having observed high CD8, T-cell chemoattractants, and ISGs in hot sarcoma tumours, we investigated if the observations in lung adenocarcinoma also held true in sarcoma subtypes. Investigating subunits of the immunoproteasome, we again observed the clearest difference between hot and cold UPS (Fig. 6c). The difference between hot and cold was least clear in DDLS (Fig. 6c). No clear difference in constitutive proteasomal subunit expression was observed between immune-hot and -cold in any sarcoma subtype (Supplementary Figure 7). Preclinical data indicate immunoproteasome subunit expression is higher in immune cells than tumours cells, and that radiotherapy can increase subunit expression in both populations (Supplementary Figure 7).

In total, these analyses provide clinical evidence that, in patients who progress after radiotherapy, immune-hot UPS is a prime candidate for 4-1BB-targeting immunotherapies. In addition, hot UPS and other immune-hot sarcoma subtypes share many of the metrics predictive of positive outcomes to immunotherapy in established ICI-responsive cancers.

## Discussion

Relapse with systemic metastatic disease after successful treatment of primary disease with neo-adjuvant radiotherapy and surgery is the major cause of disease related mortality in patients with sarcoma ^2,4^. Immunotherapy is having a transformative impact in this regard in a number of cancers ^6,7,45^. Data from the SARC028 trial have shown encouraging activity of pembrolizumab in patients with UPS and DDLS, but poor response rates in synovial sarcoma, leiomyosarcoma and osteosarcoma ^8^. The use of immunotherapy in sarcoma patients is beset with the same issues that arise with other tumour types, namely patient stratification and which alternative options to use in those refractory to anti-PD-1- and/or anti-CTLA-4-based regimens.

Immune profiling in the sarcoma TCGA study ^14^ identified a subset of DDLS, LMS, MFS and UPS tumours with high immune infiltrates. Another early study indicated greater PD-L1 and T-cell clonality in UPS ^46^. Two landmark translational studies have now been published on the SARC028 trial of pembrolizumab, focused on UPS and DDLS. Baseline tumour biopsies indicated responders had higher infiltration of T lymphocytes and a higher percentage of PD-L1^+^ macrophages, as well as effector memory and regulatory T cells ^15^. Petitprez et al. carried out analyses across a number of STS cohorts, identifying five sarcoma immune classes with histological subtypes evenly distributed across the most immune rich classes. Patients in the SARC028 trial with high B cells and tertiary lymphoid structures (TLSs) exhibited the highest objective response rate to PD-1 blockade ^12^. No objective responses were observed in the two immune-low classes, with most LMS tumours falling under the ‘immune desert’ class. Our use of a binary classification method is consistent with the consensus above, with hot tumours found distributed across UPS, MFS and DDLS, with immune-hot LMS tumours less frequent. This binary approach has revealed a number of new insights due to the ability to compare across sarcoma subtypes, and to immunotherapy-responsive cancers.

In regards to CD8, CD4 and Treg estimates, hot tumours across all four sarcoma subtypes were broadly similar to hot tumours in immunotherapy responsive cancers. Hot LMS, whilst significantly less prevalent than hot tumours in UPS, MFS, or DDLS is not an outlier in regards to the pattern of immune infiltration or the correlation of CD8, Treg and CD19 levels observed in 4-1BB-high tumours. The SARC028 trial and SU2C-SARC032 (phase II trial of neoadjuvant pembrolizumab and radiotherapy) have focused on UPS and DDLS. Although patients with LMS might benefit from immunotherapy, these data suggest that their inclusion in immunotherapy trials should be driven by selection for those with immune-hot tumours. Even at approximately 30%, hot tumours occur in patients with UPS/MFS/DDLS at only half the rate seen with some immunotherapy-responsive cancers, highlighting the importance of accurate patient stratification for immunotherapy studies in sarcoma. The binary hot/cold classification used here correlates closely to CD8 estimates. The CD3+ and CD8+ approach of Immunoscore ^47^ represents an approach that approximately reproduces the hot/cold classification method in a clinical setting. The recently published data on TLS ^12^ supports the potential benefit of including a B cell marker as a means of improving the performance of a predictive classifier.

Translational work on SARC028 ^12,15^ is reflective of findings in other cancers where many patients fail to respond to PD-1 blockade alone. Other tumour types have benefitted from dual checkpoint blockade ^6^ and data from the Alliance A091401 study of nivolumab and ipilimumab suggests this may also be true for sarcoma ^9^. Agents capable of expanding the number of patients responsive to PD-1 blockade are urgently needed. Alongside the potential to combine PD-1 blockade with radiotherapy, the findings from this study indicate 4-1BB agonists may be particularly worthy of evaluation in patients with immune-hot UPS.

Surveillance of ruptured radiation-induced micronuclei drives type-I IFN signalling in the tumour microenvironment ^16^. This type-I IFN also drives immunosuppressive signalling, such as PD-L1 ^20–22^ and has been linked both to resistance to DNA-damaging agents clinically ^42^ and in ICI-radiotherapy combinations in mouse models ^43^. We observed a higher ISG signature in hot vs cold UPS/MFS, but significantly less so in hot vs cold LMS/DDLS. It remains to be seen what impact these variables may have on radiotherapy and immunotherapy combinations. Recent preclinical data in sarcoma that used primary vs transplanted tumour indicated basal and post-radiotherapy ISG signatures were similar, yet only transplanted tumours could be cured by radiotherapy plus anti-PD-1 ^48^. This indicates that overcoming other factors such as immune tolerance may be of greater concern for clinical responses.

The neoadjuvant pembrolizumab and radiotherapy SU2C-SARC032 trial ^49^ is restricted to UPS or dedifferentiated/pleomorphic LPS of the extremity without metastatic disease. Translational work from that trial, as for SARC028, is likely to advance our understanding of responses to PD-1 checkpoint blockade plus radiotherapy in sarcoma. Preclinical data have shown the possible deleterious effects of radiotherapy on non-tumour lymphoid tissue ^50,51^. With the identification of TLS in sarcoma and the link to responsiveness to pembrolizumab ^12^, clinical data on the role of these TLS in radiotherapy plus ICI trials would be highly informative.

Maximising responses to radiotherapy is likely to require a range of immunotherapy options, from ICIs such as anti-PD-1 to co-stimulatory agonists such as 4-1BB. In the previously mentioned primary versus transplant preclinical model of sarcoma, the immune tolerance observed for primary tumours extended to resistance to radiotherapy and dual checkpoint blockade with PD-1 and CTLA-4 ^48^. Single-cell sequencing revealed low numbers of activated T cells in primary tumours even after anti-PD-1 treatment. Co-stimulatory agonists may represent an approach to overcome this immune tolerance ^31^. 4-1BB (CD137) is an inducible co-stimulatory receptor that, when activated, has been shown to be anti-apoptotic and restore effector function in dysfunctional T-cells ^30^. The natural ligand of 4-1BB (CD137L/4-1BBL) is expressed on DC, macrophages and B cells. Our data show that UPS tumours are amongst the highest expressors for 4-1BB and 4-1BBL, potentially pointing the way towards development of 4-1BB-targeted agents in these diseases. Clinical development of 4-1BB agonists has, so far, been hampered by weak efficacy in the case of utomilumab ^52,53^, or liver toxicity for urelumab that has restrained dosing below levels where efficacy has been observed ^54,55^. There are significant functional differences between each agent, with the IgG4 urelumab shown to increase 4-1BBL-dependent clustering of 4-1BB, in contrast to the ligand-blocking binding site of the IgG2 utomilumab ^37^. The presence of both 4-1BB and 4-1BBL in hot UPS marks it as a prime candidate for a non-ligand blocking agonist.

Clinical progress with 4-1BB-targeted agents has been impacted by the complex interplay between ligand blocking, Fab region binding, and Fc receptor (FcR) crosslinking in determining both agonist activity and hepatic toxicity. Crosslinking by FcγRIIB expressed on liver sinusoidal endothelial cells has been implicated as a likely cause of urelumab-induced toxicity ^56,57^. Studies have focused on the development of bispecific agonists promoting agonist localisation to the tumour microenvironment ^56,58^, or have sought to understand the relative roles of Fab region binding versus FcR clustering. New preclinical studies of 4-1BB agonists have shown that strong anti-tumor efficacy can be generated whilst avoiding both 4-1BBL ligand-blocking and liver toxicity due to crosslinking ^57,59^. One study has indicted that 4-1BB antibodies that are weak agonists from Fab binding alone, but require inhibitory Fc-receptor FcγRIIB-mediated crosslinking, retain anti-tumour activity in the absence of liver toxicity ^57^. The agonist described in that study, LVGN6051, has now entered clinical trials as a monotherapy and in combination with anti-PD-1 (NCT04130542).

The complexity of how best to target 4-1BB has been further demonstrated by studies showing it to be highly expressed on intratumoural versus peripheral Tregs ^31,32^. Preclinical studies indicate the therapeutic potential of targeting 4-1BB with isotypes promoting antibody-dependent cellular cytotoxicity (ADCC)-mediated depletion of intratumoral Tregs ^31,32^. Our correlation analysis indicates CD8 and Tregs both correspond to high 4-1BB levels, but more granular data shows a high degree of heterogeneity when examining CD19, CD8 and Treg estimates together with 4-1BB expression levels. This brings us back to the critical importance of patient stratification. It is likely that the therapeutic responses to either agonist or ADCC-promoting isotypes targeting 4-1BB will be determined by the presence of specific profiles of immune populations in the tumour microenvironment of sarcoma patients.

Radiotherapy increases the number of intratumoral Tregs in preclinical studies ^20,60^ and the percentage of 4-1BB-positive Tregs but not levels of 4-1BB on CD8^+^ cells ^60^. This potentially points to 4-1BB depletion as more appropriate to future radiotherapy combinations. 4-1BB agonism has been shown to improve the therapeutic outcome of both anti-PD-1 and anti-CTLA-4 when combined preclinically with radiotherapy or cisplatin-radiotherapy ^61,62^. A bispecific 4-1BB-VEGF aptamer has also been shown to increase both efficacy of radiotherapy at the primary site and to reduce disease burden at out-of-field lung metastases ^63^. These studies show the potential of 4-1BB-targeting agents to promote both tumour control at the primary irradiated site and the distal abscopal effect. Alongside the previously mentioned LVGN6051, many other new 4-1BB agonists (ADG106, AGEN2373, ATOR-1017, PRS-343) continue to enter phase I trials. The variety of IgG types, Fab affinities, or bispecific variants will hopefully lead to clinical data to guide the best approach to targeting 4-1BB whilst reducing liver toxicity.

In summary, our data point to a group of immune-hot UPS patients that may be highly amenable to immunotherapies. These patients, who progress after radiotherapy, are strong candidates for emerging 4-1BB-targeting agents. Questions remain on the best approach to targeting 4-1BB. However, preclinical studies suggest 4-1BB-targeting agents, in combination with radiotherapy, have the potential to improve outcomes through a reduction in metastatic disease. Taken in totality with recent publications on the SARC028 study, these data indicate many more sarcoma patients could benefit from immunotherapy.

## Materials and Methods

### Retrospective STS patient samples

Archival histopathological samples were retrieved from the Royal Marsden Hospital tissue bank. This study was approved by the institutional review board (Committee for Clinical Research No. CCR4852). Consent was confirmed for all patients. Pre-treatment biopsy specimens were sourced from archival blocks from patients who received neoadjuvant radiotherapy followed by surgical resection. Archival tissue was retrieved from a further 2 UPS patients who did not receive pre-operative radiotherapy. Retrieved samples were limited to patients diagnosed with soft tissue sarcoma of the extremities with the following histological subtypes – undifferentiated pleomorphic, myxofibrosarcoma and myxoid liposarcoma. Samples from patients with recurrent or metastatic soft tissue sarcomas were excluded.

### NanoString gene expression analysis

H&E sections from FFPE samples were outlined to guide macrodissection of viable areas of tumour by an expert soft tissue sarcoma pathologist (KT). Macrodissection was performed on nuclear fast red-stained sections. RNA was extracted using the Allprep DNA/RNA FFPE kit (Qiagen) using the manufactures protocol. Extracted RNA was quantified using a NanoDrop spectrophotometer (Thermo Fisher) and quality assessed on a BioAnalyzer 2100 (Agilent). Samples were stored at −80 °C before further analysis. 100 ng RNA was used for analysis using the Human nCounter PanCancer Immune Profiling Panel (NanoString Technologies). A customised 30-gene panel was included alongside the standard probe set. Custom probes were included against the following gene transcripts; *APEX1, APOBEC3B, ATR, BATF3, BRCA1, BRCA2, CFLAR, CLEC9A, DDB2, H2AFX, IFIT3, LIG4, CGAS, MDC1, MICA, MLH1, NBN, NLRP9, OAS1, OAS2, PARP1, PCNA, RAD51, RBBP8, RPA3, STK39, STING, TRADD, TREX1, XRCC4*. Normalisation from housekeeping genes and for hybridisation was performed using nSolver v4.0.70 (NanoString). Geometric mean of negative controls was chosen for background thresholding. Geometric mean of positive controls was used to compute normalisation factor and lanes with a range outside 0.3-3 were flagged. One sample had to be excluded due to a normalisation flag, this was a UPS sample with the highest 4-1BB level. Immune cell scoring from NanoString data was performed using the method described previously ^34^. Differentially expressed genes were determined in nSolver. Foldchange cutoff of 2 is shown for transcripts significantly higher in UPS versus MLPS, with a FDR-corrected p-value of less than 0.05. Heatmaps were plotted using the ComplexHeatmap package. On log2 transformed data, hierarchical clustering was performed and k-means clustering of 2 is shown.

### Immune cell multiplex immunohistochemistry

FFPE tissue sections were stained for multiplex fluorescence-based IHC using the opal 7-colour kit (Akoya Biosciences). Sections were dewaxed and rehydrated. Sequential rounds of antigen retrieval, blocking, primary antibody incubation, wash, secondary-HRP conjugate and opal reagent incubation were performed using the recommended manufacturers’ guidelines. Primary antibodies used were: mouse anti-human CD8 (Dako, clone C8/144B, 1:800, pH 9 retrieval), mouse anti-human CD68 (Dako, clone PG-M1, 1:750, pH 6 retrieval), mouse anti-human CD20 (Dako, clone L26, 1:1000, pH 6 retrieval), mouse anti-human FOXP3 (Abcam, clone 236A/E7, 1:600, pH 6 retrieval), rabbit anti-human CD4 (Abcam, clone EPR6855, 1:1000, pH 9 retrieval). For each marker a single-stain control tissue sample was included. Multiplex IHC slides were imaged on a Vectra 3 (Akoya Biosciences). Spectral unmixing and cell phenotype identification was performed using inForm software (Akoya Biosciences). Digital immune phenotype plots were created by assembling xy data from identified cell phenotypes in individual tile scans into a whole biopsy composite using ggplot. To identify areas of localisation more easily, granularity was reduced by hexbin-derived heatmaps using the hexbin package.

### TCGA sarcoma data analysis

Transcriptomic data from the TCGA were accessed using cgdsr. TCGA sarcoma data were restricted to UPS, MFS, DDLS and LMS, as were pooled sarcoma data presented in figure 4. We maintained the separation of UPS and MFS as distinct subtypes for our analyses, although molecular data from the initial TCGA publication state that these two subtypes are not distinct and fall along a spectrum ^14^. TCGA data for other cancer types were used as labelled and not restricted. Where it was judged to aid the clarity of presentation, data were log2 transformed. For survival probability analysis, 61 samples received radiotherapy (2 DDLS, 17 LMS, 17 MFS, 25 UPS), with 128 receiving no radiotherapy (32 DDLS, 68 LMS, 8 MFS, 20 UPS). Samples where radiotherapy status could not be clearly determined were excluded. Cancers in the TCGA dataset were selected for comparative purposes to sarcoma due to their responsiveness to immunotherapy, or those commonly referred to as immune-hot, immune-cold or with low mutational burden.

### MethylCIBERSORT derived hot/cold classification and immune cell composition

Use of MethylCIBERSORT to generate binary ‘immune-hot’ and ‘immune-cold’ subgroups and immune cell population estimates has previously been published and is described in detail^13^. Correlation analyses of estimated immune populations based on 4-1BB status were performed using the R packages cor and corrplot.

### Code and data availability

The data accessed in this study are available from The Cancer Genome Atlas Project. The code used for analyses and non-TCGA data are available from the authors upon reasonable request. Code and data availability for immune composition and hot/cold classification has been previously outlined ^13^.

## Acknowledgements

This study was supported by RM/ICR NIHR Biomedical Research Centre (MJM, HGS, KJH), Rosetrees Trust (KJH, grant numbers M48 and M444), Cancer Research United Kingdom (DM, AAM, KJH) and Anthony Long Charitable Trust (KJH). We would like to acknowledge and thank Priya L. Narayanan and Nick Trahearn from the Yuan lab at The Institute of Cancer Research for digital spatial composite assistance and the Breast Cancer Now pathology core facility.

---

**Supplementary Table 1:** Patient characteristics of soft tissue sarcoma patients. Cohort of n=27 soft tissue sarcoma patients of which n=25 received neoadjuvant-radiotherapy and n=2 received no radiotherapy.

| **Histology, n** |  |
| --- | --- |
| Undifferentiated pleomorphic sarcoma (UPS) | 11 |
| Myxoid liposarcoma (MLPS) | 10 |
| Myxofibrosarcoma (MFS) | 6 |
| **Grade, n** |  |
| 1 | 4 |
| 2 | 10 |
| 3 | 8 |
| Not known | 5 |
| **Site, n** |  |
| Buttock | 4 |
| Thigh | 17 |
| Calf | 1 |
| Shoulder girdle | 5 |
| **Depth, n** |  |
| Superficial | 2 |
| Deep | 25 |
| **Size, n** |  |
| <5 cm | 2 |
| 5-10 cm | 12 |
| 10-15 cm | 6 |
| >15 cm | 7 |
| **Local or Distant Progression, n** |  |
| Local | 2 |
| Distant | 10 |
| NA/unknown | 15 |

**Supplementary Figure S1.**
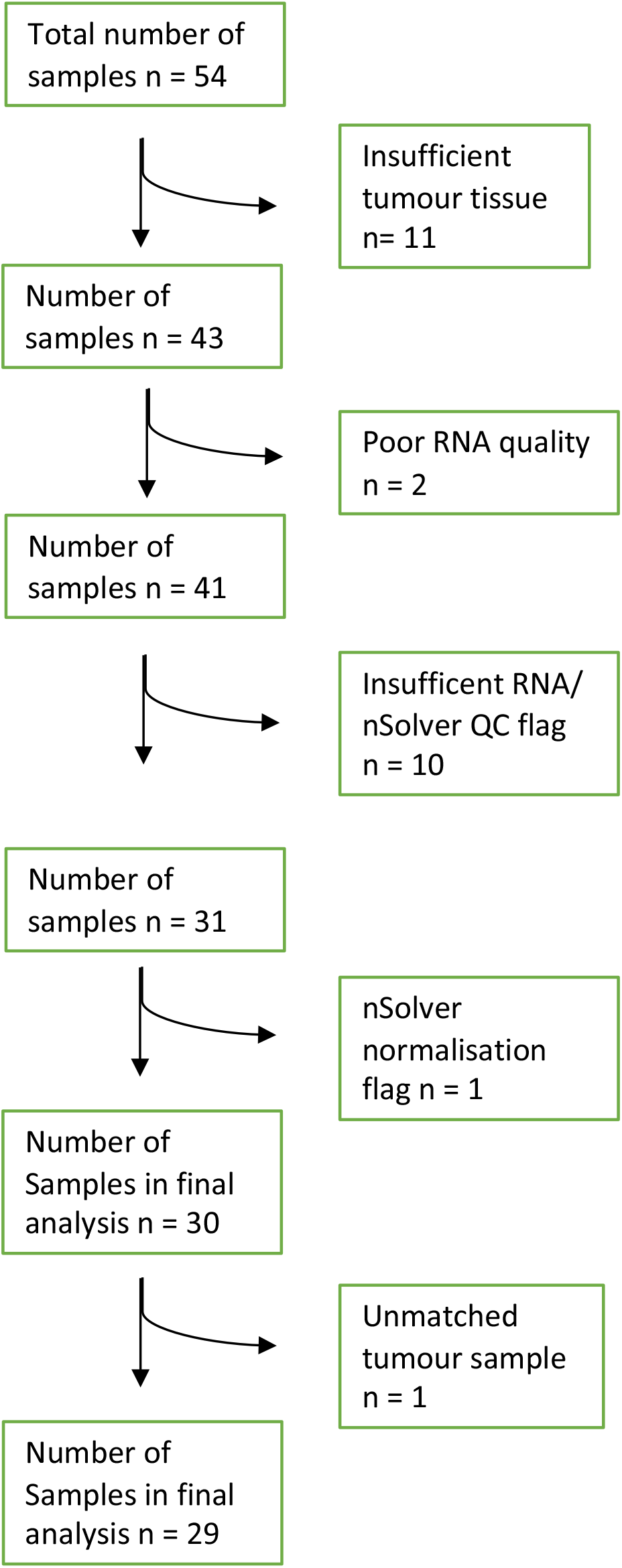
Chart showing sample loss due to quality control for Nanostring analysis shown in Figure 1. Initial retrospective samples requested were from n=27 patients with paired pre-radiotherapy biopsies and surgical resections resulting in n=54 FFPE samples in total. Flow chart indicating loss of samples from quality control (QC) at each step. Pathology outlines for sufficient tumour, RNA quality, Nanostring/nSolver QC flag (FFPE degradation of RNA too high), a single normalization error, a sample where only the surgical resection and not the pre-treatment biopsy was available. Of the final n=29 samples, n=21 were pre-treatment biopsies and n=8 were from subsequent surgical resections. Pretreatment biopsies corresponded to MLPS n=9, UPS n=7, MFS n=5. From surgical resection samples, as only n=6 were samples paired with pre-treatment biopsies, these were not investigated further.

**Supplementary Figure S2.**
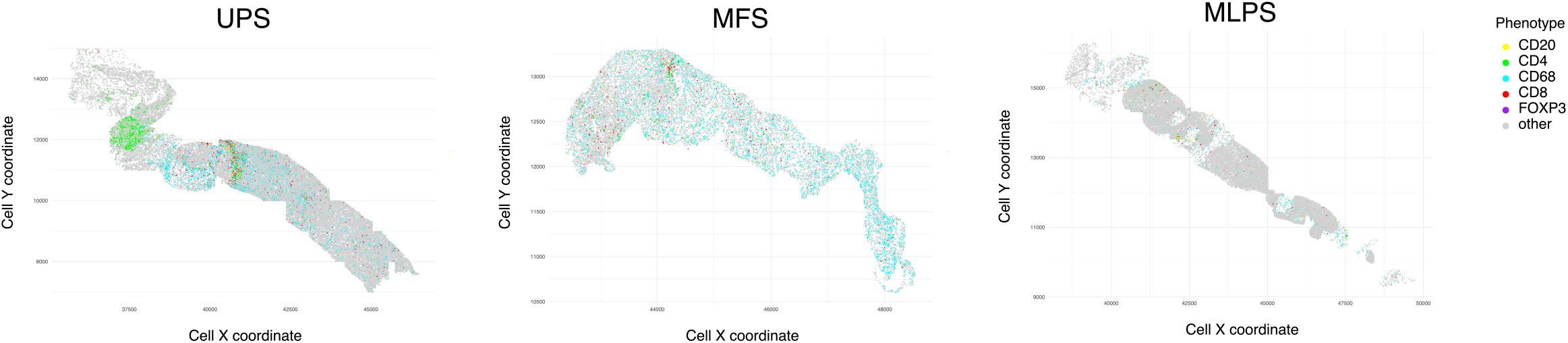
Multiplex immunohistochemistry digital spatial plots. Spatial plots of xy and cell phenotype data derived from Vectra multiplex IHC imaging and cell type identification. Plots show all populations identified, and correspond to the cell density plots for individual populations shown in figure 2c.

**Supplementary Figure S3.**
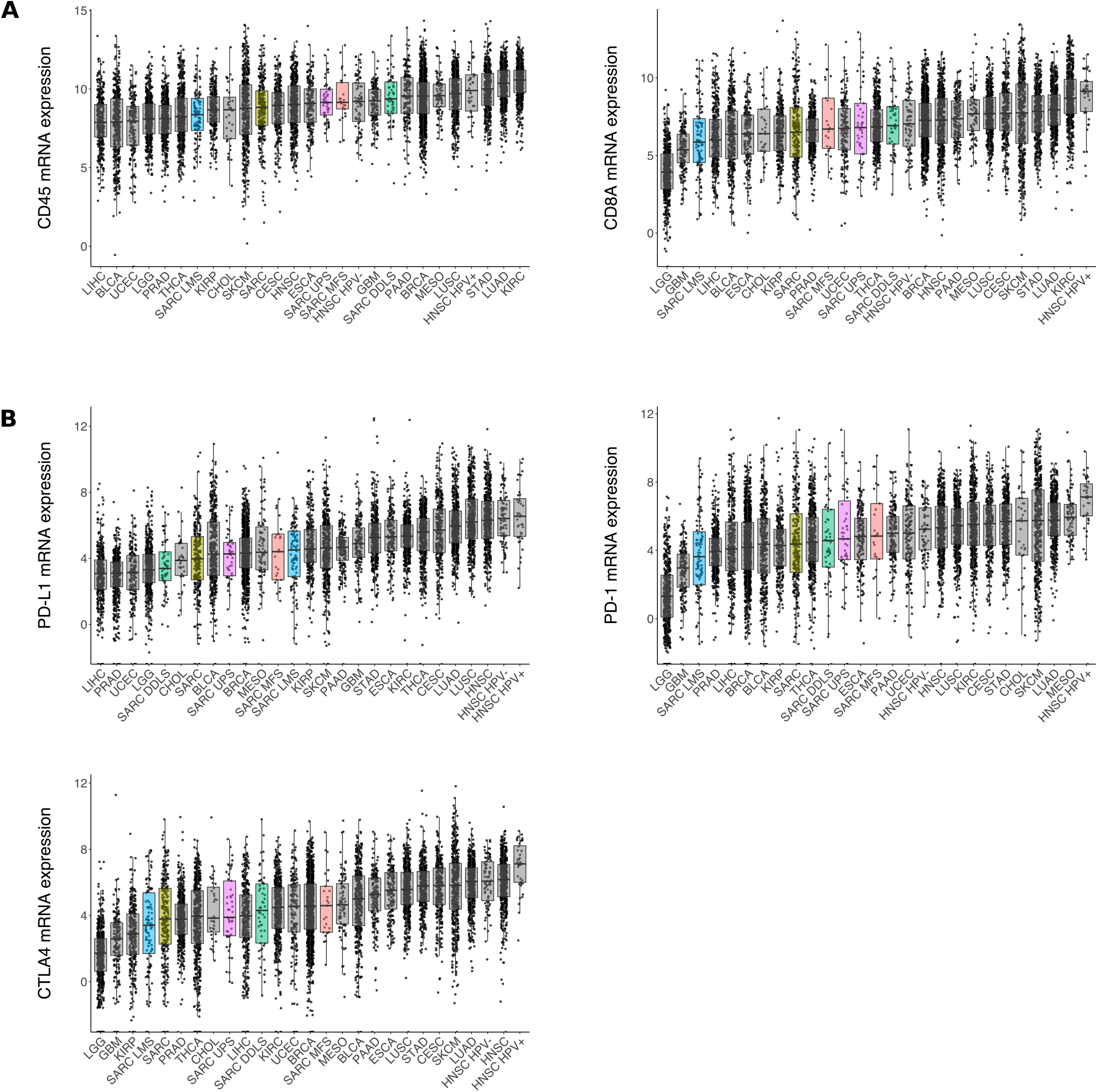
TCGA expression data for selected immune cell population transcripts and current clinical immunotherapy targets. Expression data for sarcoma and split sarcoma subtypes compared to selected other cancer types in the TCGA dataset. **a.** *PTPRC* (CD45) and *CD8A*, **b.** *CD274* (PD-L1), *PDCD1* (PD-1) and *CTLA4*.

**Supplementary Figure S4.**
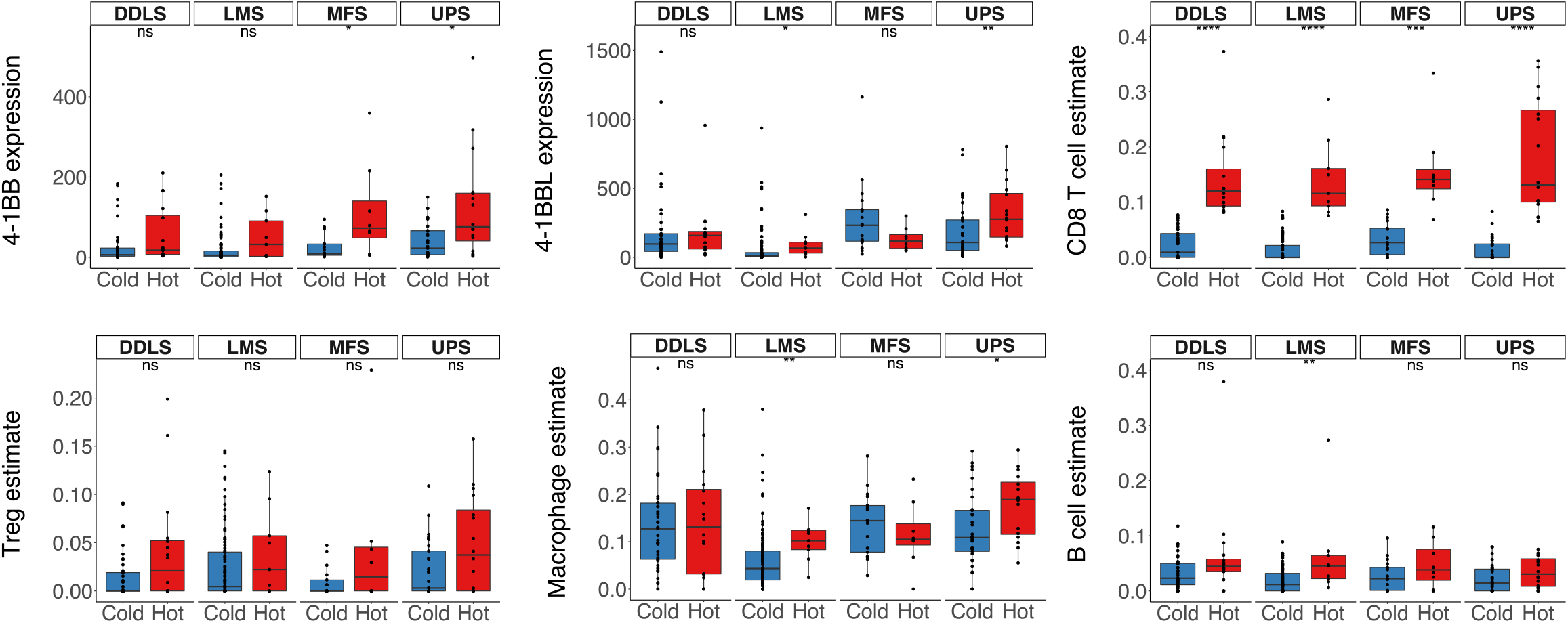
Statistical comparison between hot and cold populations by sarcoma subtype corresponding to Figure 4. Data from Figure 4 showing statistical comparisons between hot and cold populations for each sarcoma subtype for *TNFRSF9* (4-1BB) and *TNFSF9* (4-1BBL) mRNA expression, as well as CD8, Treg, monocyte/macrophage and B cell population estimates. Significance shown t-test, *p<0.05, *p<0.01., *p<0.001., *p<0.0001.

**Supplementary Figure S5.**
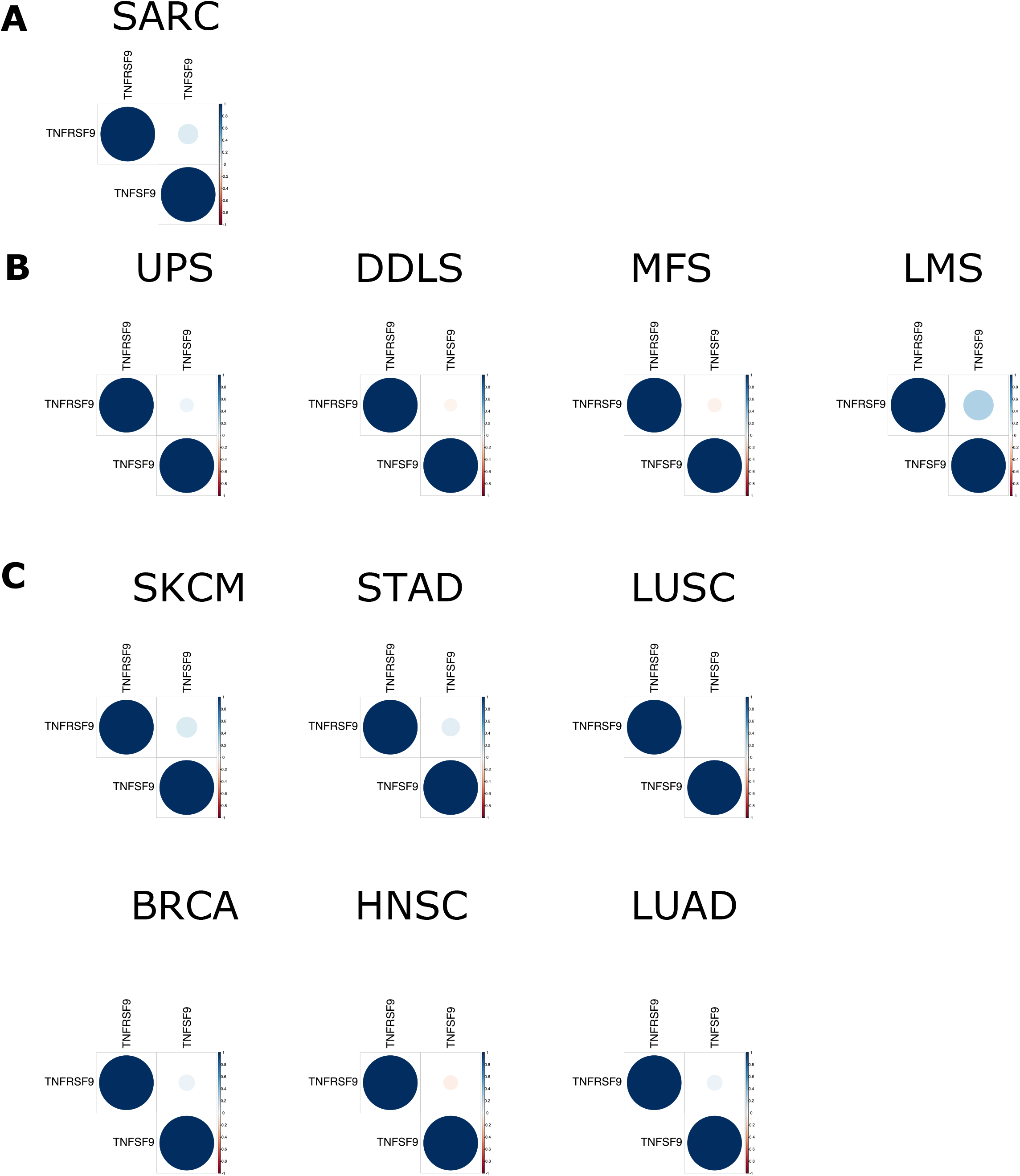
Correlation analysis does not indicate a link between *TNFRSF9* (4-1BB) and *TNFSF9* (4-1BBL) expression levels. Pearson’s correlation analysis was performed for TCGA samples on **a.** all sarcoma samples, **b.** sarcoma samples split by subtype and **c.** for selected immunotherapy responsive cancers.

**Supplementary Figure S6.**
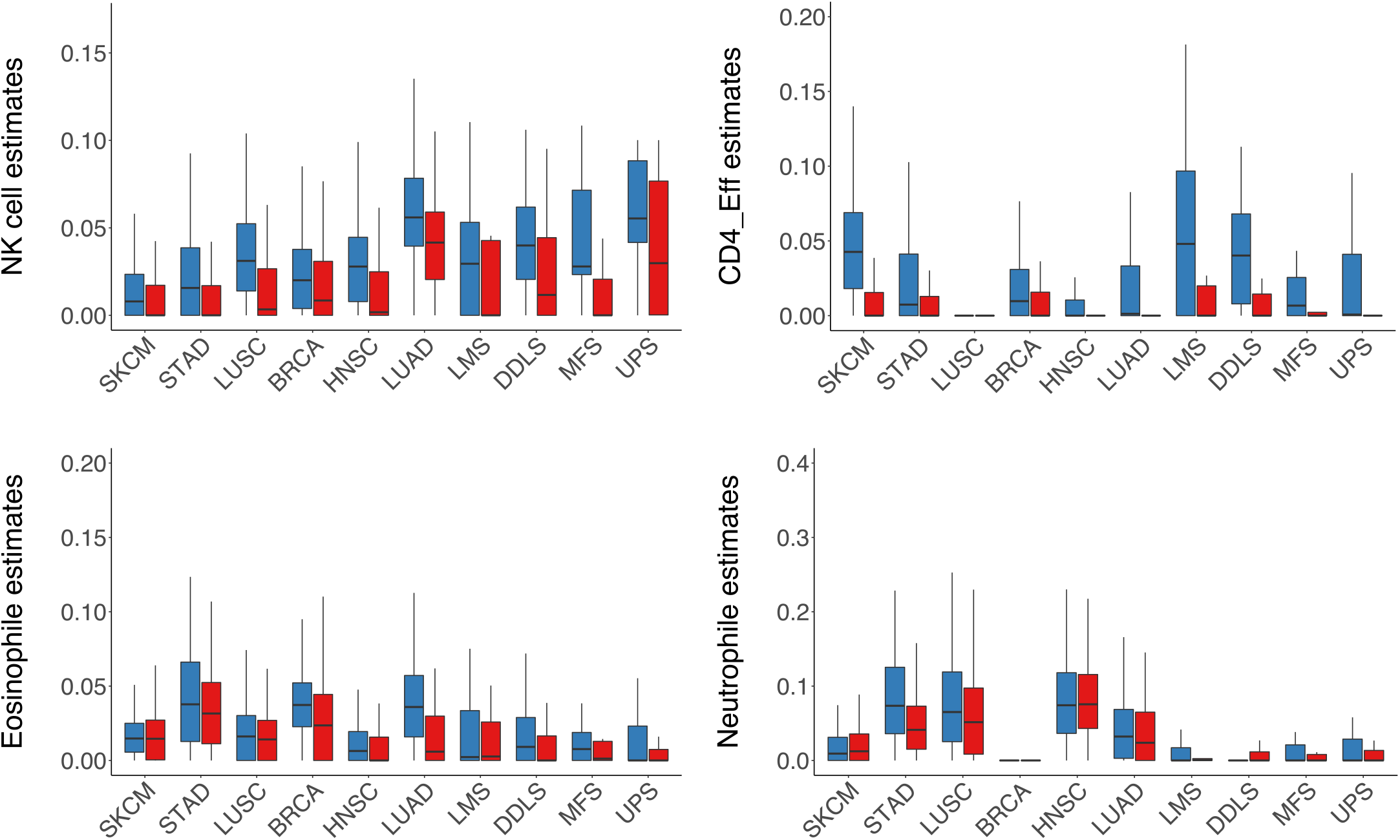
MethylCIBERSORT derived population estimates for additional immune cell populations. Immune population estimates are displayed as shown in figure 4g for NK cells, CD4 effector cells, eosinophils and neutrophils. Comparison between sarcoma subtypes and immunotherapy responsive cancers. Split by hot (red) and cold (blue) MethylCIBERSORT derived binary immune classification status.

**Supplementary Figure S7.**
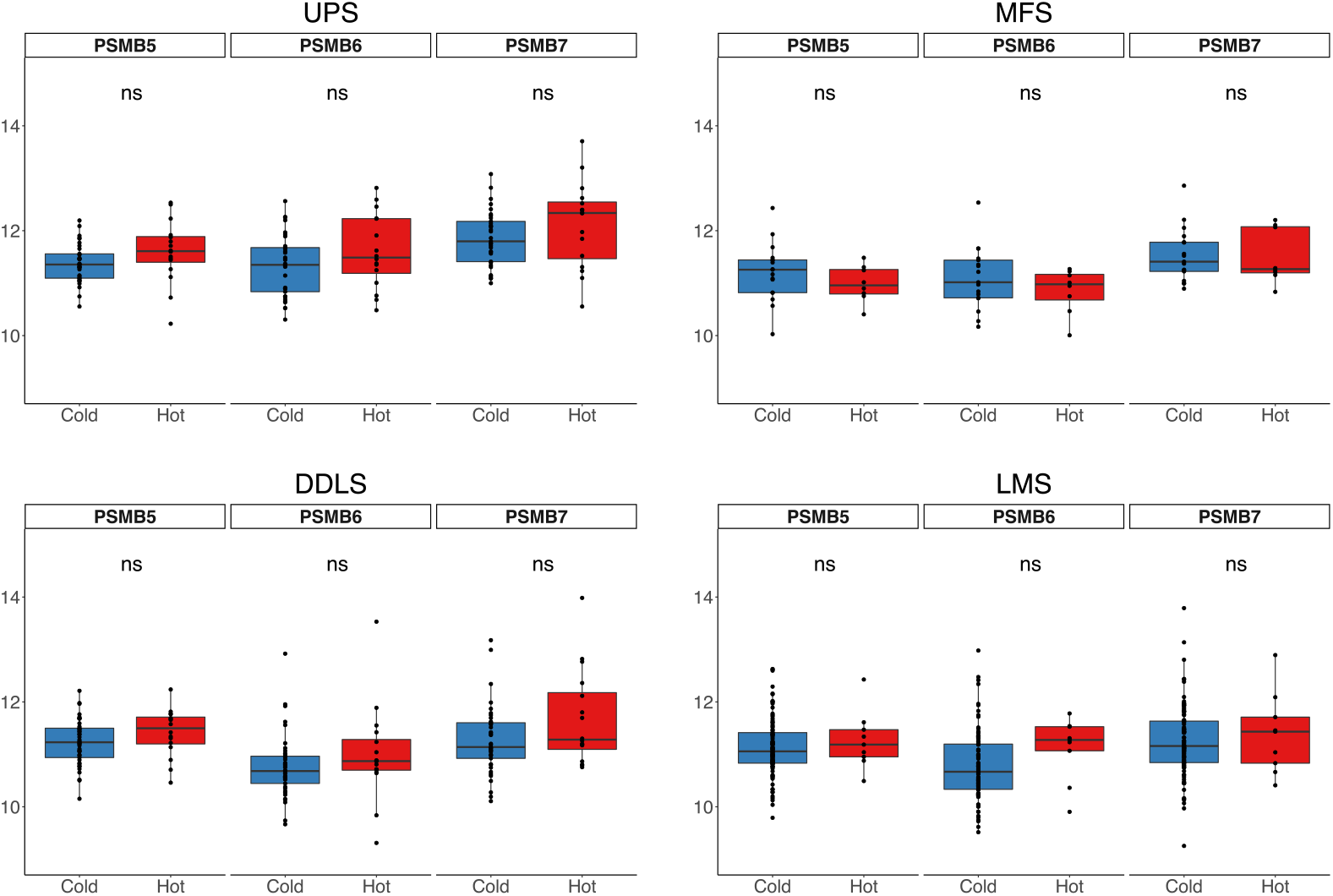
Expression of constitutive proteomic subunits does not differ based on binary immune classification status. The expression of the constitutive proteasomal subunits *PSMB5*, *PSMB6* and *PSMB7* was assessed in TCGA data for each sarcoma subtype. Statistical comparisons were carried out by t-test between hot (red) and cold (blue) MethylCIBERSORT derived binary immune classification status.

